# Household triclosan and triclocarban exposure impacts the adult intestinal microbiome but not the infant intestinal microbiome

**DOI:** 10.1101/126334

**Authors:** Jessica V. Ribado, Catherine Ley, Thomas D. Haggerty, Ekaterina Tkachenko, Ami S. Bhatt, Julie Parsonnet

## Abstract

In 2016, the US Food and Drug Administration banned the use of specific microbicides in some household and personal wash products. This decision was due to concerns that these chemicals might induce antibiotic resistance or disrupt human microbial communities. Triclosan and triclocarban (referred to as TCs) are the most common antimicrobials in household and personal care products, but the extent to which TC exposure perturbs microbial communities in humans, particularly during infant development, was unknown. We conducted a randomized intervention of TC-containing household and personal care products during the first year following birth to characterize whether TC exposure from wash products perturbs microbial communities in mothers and their infants. Longitudinal survey of the intestinal microbiota using 16S ribosomal RNA amplicon sequencing showed that TC exposure from wash products did not induce global reconstruction of either infant or maternal intestinal microbiotas following 10 months of exposure after birth. However, broadly antibiotic-resistant species from the phylum Proteobacteria were enriched in stool samples from mothers in TC households only after the introduction of triclosan-containing toothpaste. Despite the minimal effects of TC exposure from wash products on the gut microbial community of infants and adults, these results suggest detected taxonomic differences are associated with potential harmful effects on host physiology, highlighting the need for consumer safety testing of self-care products not subject to the ban on the human microbiome and health outcomes.

## Introduction

Triclosan and triclocarban (TCs) are chlorinated, broad-spectrum antimicrobial chemicals found in thousands of consumer and industrial products. They are present most notably in personal wash products including toothpaste and liquid soaps (triclosan) and bar soaps (triclocarban). In 2016, a Food and Drug Administration (FDA) ruling banned the use of TCs and 17 other antimicrobial chemicals in over-the-counter wash products, driven by the concern that the use of these products contributed to antibiotic resistance and might negatively affect human health, either through endocrine disruption or modification of the human microbiota(*1*). Notably, many other TC-containing products, such as toothpaste, fabrics and plastic goods (including toys), were not subjected to the ban.

To date, limited data exists regarding the effects of TCs on the human microbiota(*2*). The microbes that occupy the human body in niches from the gut to the skin have diverse roles in human health, ranging from metabolic support to immunomodulation. Imbalances in these microbial communities are implicated in a wide variety of diseases(*3*). The extent to which triclosan exposure may induce microbial perturbations has been studied in fish and rodent models with conflicting outcomes (reviewed in (*4*)). Triclosan exposure restructures the juvenile fish microbiome(*5*), but results in recoverable alterations following short-term perturbation in adult fish(*6*). Adolescent rats receiving oral triclosan at levels comparable to human exposures develop lower microbial diversity in the gut and more prominent changes in taxonomic composition than in adult rats(*7*). While triclocarban exposures are less studied, in pregnant rats and their offspring less than 10 days old, exposure lead to lowered phylogenetic diversity and revealed a dominance of the Proteobacteria phylum in the gut(*8*). In a small, randomized cross-over human study, TC wash product exposure did not induce major perturbations of the oral and gut microbiomes(*9*). This finding supports other studies that have shown minimal impact of triclosan on dental microbial ecology, despite slowing the progression of periodontitis(*10*).

The core microbiome of humans is established in the first few years of life(*11*). Disruptions to the microbiota early in development by extrinsic factors, such as antibiotics, can have long-term impacts on metabolic regulation(*12*) and can delay normal microbiota maturation(*13*). The impact of TC exposure through household and personal care products on the developing microbiota is unknown. As a nested, randomized intervention within Stanford’s Outcomes Research in Kids (STORK), a prospective cohort study of healthy mothers and infants(*14*), we provided commercially available wash products containing or not containing TCs (TC and nTC arms, respectively) to evaluate their relative impact on the maternal and infant intestinal microbiota over the first year of the infant’s life.

## Results

### Study Demographics

Thirty-nine households from the STORK cohort met our inclusion criteria (i.e., at least 5 of 6 expected stool samples available from the household). Complete sampling for both infants and mothers for three time points after birth was available for 26 households, and one sample was missing for 13 households. Home visits and sample collection occurred on average 74 (14-124), 200 (135-256) and 317 (241-377) days following birth (Supplementary figure 1). These days correspond to approximately 2.7, 6.6, and 10.6 months, referred to as 2, 6, and 10 months hereafter. The average age of mothers in this subset was 34 years and 46% were of Hispanic origin (Table 1).

**Table 1:** Selected characteristics of study sample (N=39 households). Demographics are self-reported at the time of enrollment. Age, individuals in the household, and the sample collection windows are reported as the median and range. Sample collection times are relative to the infant birth date. Not applicable for cleaning products at work indicates the mothers were unemployed at the time, and not receiving additional TC exposure similar to mothers that do not work with cleaning products.

|  | **TC**<br>**(N=17)** | **nTC**<br>**(N=22)** |
| --- | --- | --- |
| **Individuals residing in household** | 4 (3-10) | 3 (2-11) |
| **Maternal age (years)** | 33 (26-41) | 34 (28-42) |
| **Ethnicity** |  |  |
| Hispanic | 11 (64.7%) | 7 (31.8%) |
| Non-Hispanic | 6 (35.3%) | 15 (68.2%) |
| **Bathing habits** |  |  |
| Less than daily | 5 (29.4%) | 3 (13.6%) |
| At least daily | 12 (70.6%) | 19 (86.4%) |
| **Pets** |  |  |
| Yes | 6 (35.3%) | 5 (22.7%) |
| No | 11 (64.7%) | 17 (77.3%) |
| **Mother's use of cleaning products at work** |  |  |
| Yes | 5 (29.4%) | 6 (27.3%) |
| No | 5 (29.4%) | 11 (50.0%) |
| Not applicable | 7 (41.2%) | 5 (22.3%) |
| **Days after delivery for each sample collection window** |  |  |
| 2 months | 88 (14-124) | 74 (27-122) |
| 6 months | 199 (135-256) | 199 (146-255) |
| 10 months | 330 (259-375) | 319 (241-277) |

### Randomization to TC-containing household and personal products is sufficient to increase triclosan exposure after 6 months

Mothers in TC households had higher spot urinary triclosan levels at 6 months when compared to those in nTC households, with a median triclosan measurement of 837.05 pg/μL compared to 76 pg/μL (p < 0.001). With one exception, levels in children were uniformly low, with a median of 38.3 pg/μL in TC households and 10.05 pg/μL in nTC households (p=0.06, Figure 1, Supplementary table 2).

**Figure 1:**
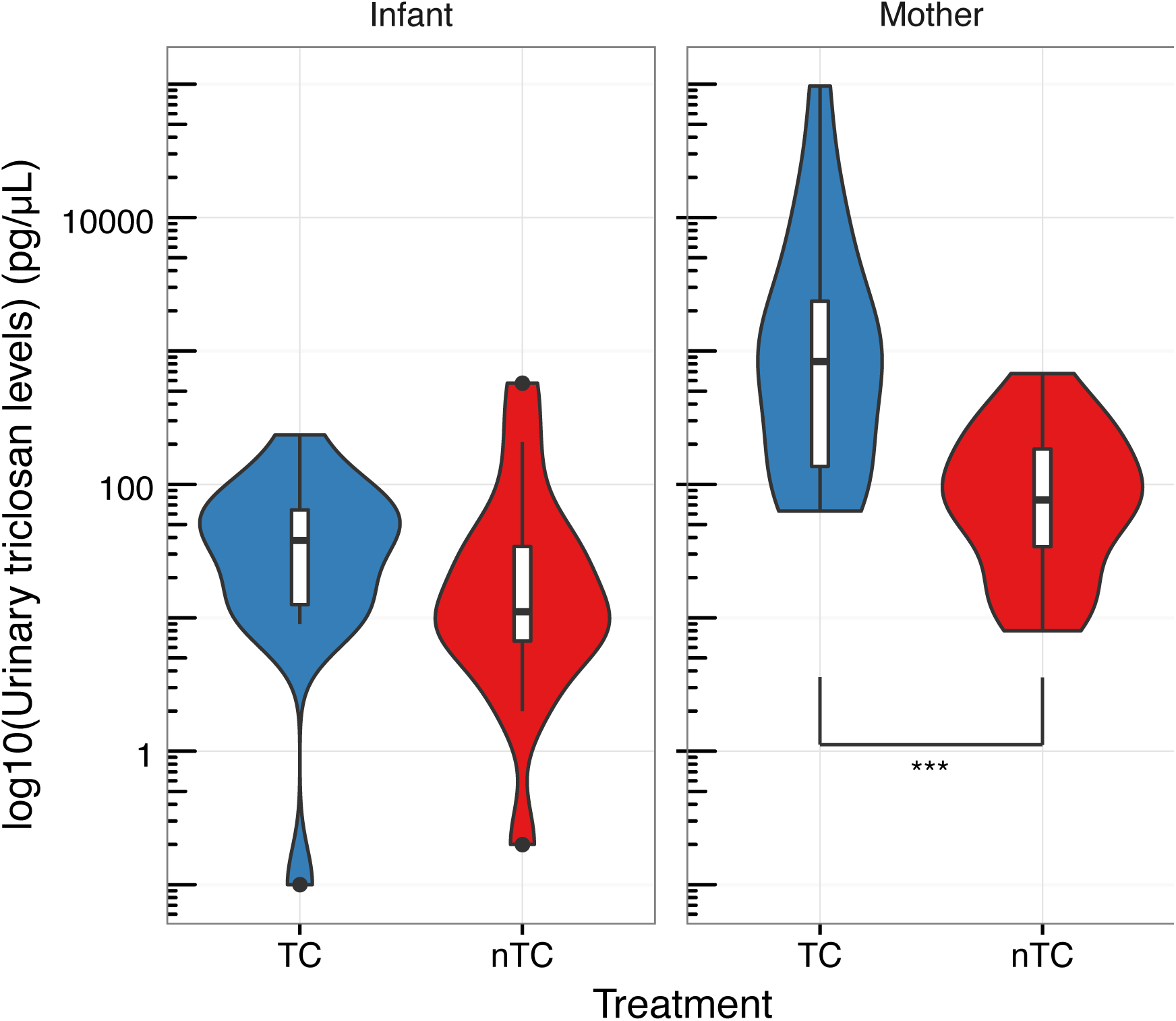
Urinary triclosan levels are elevated in TC mothers following 6 months of exposure. Urinary triclosan measurements are available for 38 mothers (17 TC, 21 nTC) and 33 infants (15 TC, 18 nTC). Mothers in TC households have statistically significant higher triclosan levels than nTC mothers (Mann-Whitney U-test, p < 0.0001). Infant TC levels between treatments are not significant (Mann-Whitney U-test, p=0.06).

### Mother and infants have distinct microbiome compositions not driven by randomization to TC-containing products

A principle coordinate analysis (PCoA) was performed to identify variability between the taxonomic structure of the samples. PCoA showed that samples segregated primarily by age, with 31.9% of variation among samples explained by the first two axes (Figure 2A). By 10 months of age, infant samples clustered more closely to the mothers’ samples (Figure 2B). Individual samples from TC and nTC households were evenly dispersed through the axes for both infants and mothers. Taxonomic classifications suggest that samples were generally similar at the phylum level among infants and among mothers, regardless of treatment arm (Supplementary figure 3). Variations between infants were not driven by factors known to influence microbial colonization, such as delivery method, breast feeding, and pets in the household (Supplementary figure 4). Maternal samples from the various time points cluster more closely by individual than by time point (Supplementary figure 5A), and we did not observe major variations in taxonomic structure between maternal samples based on TC exposure using PCoA (Supplementary figure 5B, 5C). Statistical comparisons of treatment arms with permutational multivariate analyses showed no significant association between TC exposure and microbiome composition for infants (p=0.14), but did demonstrate a significant association between TC exposure and microbiome composition in the mothers (p < 0.002).

**Figure 2:**
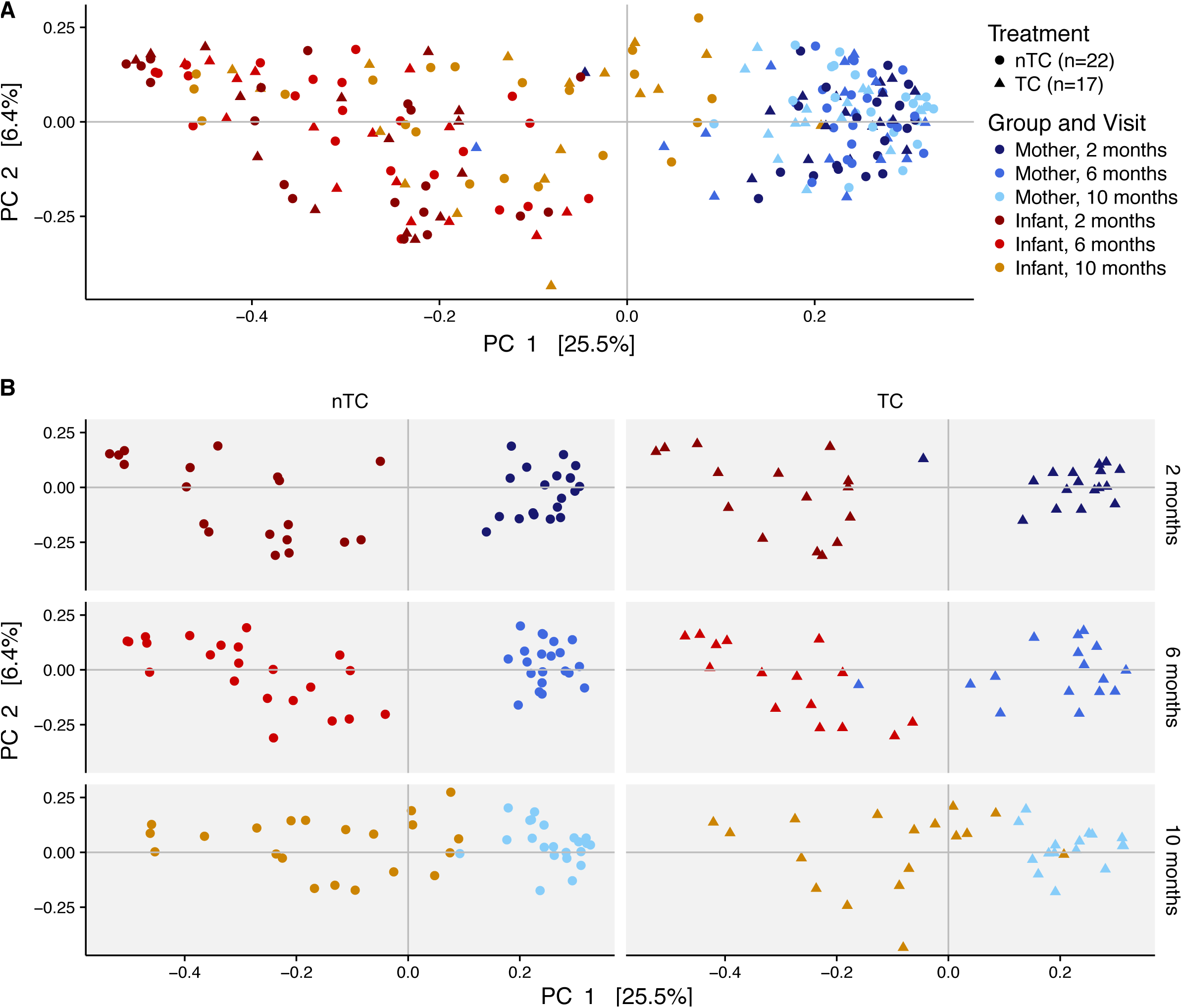
Mother and infants have distinct microbiome compositions not driven by household TC exposure. (A) PCoA of Bray-Curtis dissimilarity for all (n=221) samples shows that gut communities cluster by mothers and infants. (B) PCoA separated by time and treatment.

### Randomization to TC-containing products does not decrease gut microbial diversity in infants or mothers

Randomization to the TC arm was not associated with decreased gut microbiota diversity for infants or mothers at any of the 2, 6, or 10-month visits after infant birth (Figure 3). Specifically, diversity was not decreased in infants randomized to TC-containing products (p-values for 2, 6, and 10 months: 0.66, 0.84, 0.49). As expected, microbial diversity increased as the infants progressed through the first year of life (p < 0.001)(*15*), and this effect was not altered by randomization to TC. Diversity was not statistically significantly decreased in maternal samples randomized to TC-containing products (Mann-Whitney U-test p-values for 2, 6, and 10 months: 0.73, 0.28, 0.20), however trended to a decrease in stool diversity at the 10-month visit.

**Figure 3:**
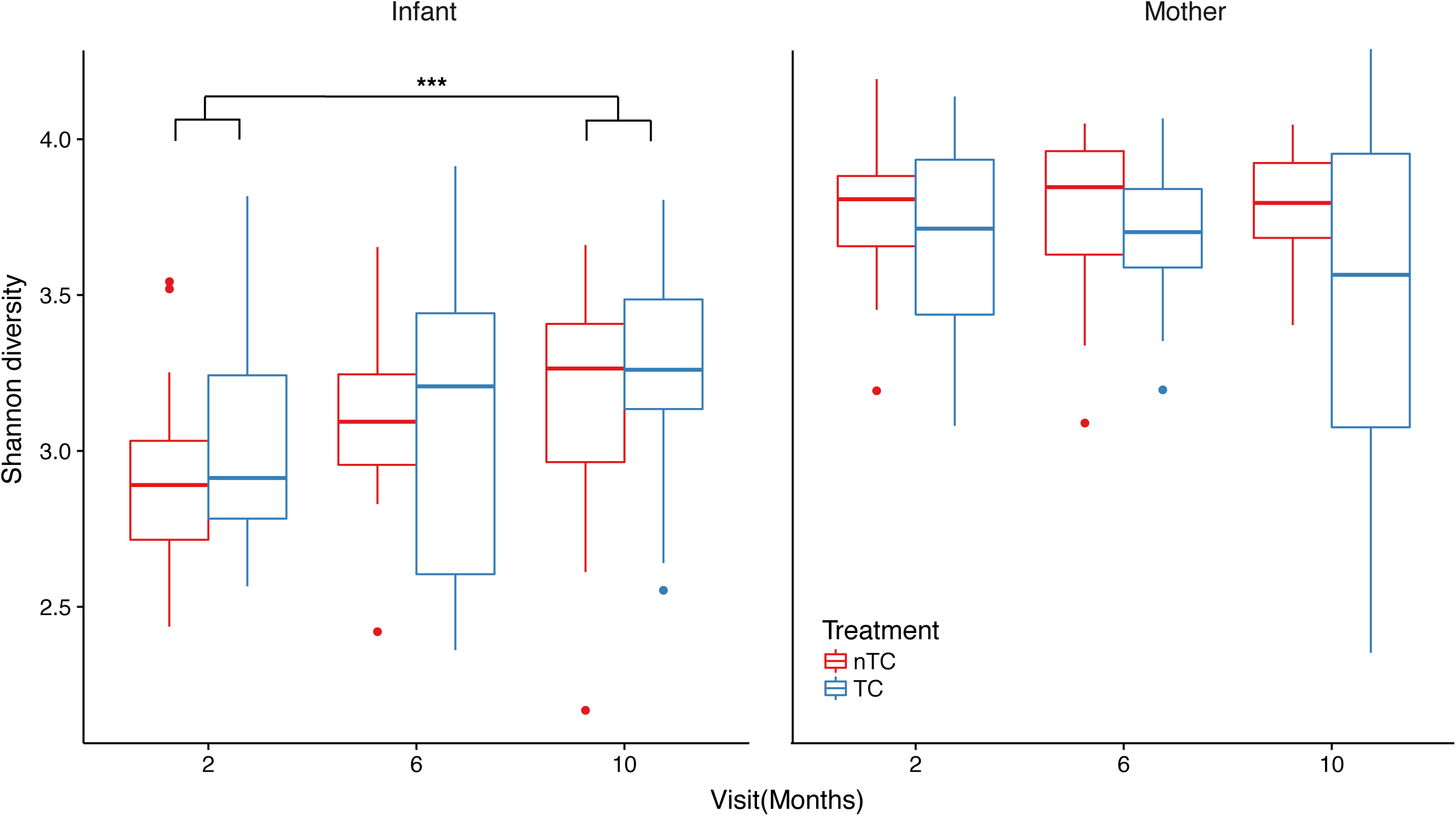
TC randomization does not decrease gut microbial diversity in infants or mothers. Shannon diversity measures are plotted as interquartile range with median for each TC exposure class and time point grouping for infants and mothers. Diversity is not decreased by exposure to TC containing products in infants (Mann-Whitney U-test p-values for 2, 6, and 10 months: 0.656, 0.842, 0.486). The combined infant cohort shows an increase in diversity as colonization occurs through the first year (Mann-Whitney U-test for 2 to 12 months, p-value < 0.001). Diversity is not decreased by exposure to TC-containing products in mothers (Mann-Whitney U-test p-values for 2, 6, and 10 months: 0.729, 0.280, 0.201).

### Intestinal exposure to triclosan through toothpaste, rather than wash products, is associated with Proteobacteria enrichment in TC households

Given this trend toward decreased diversity with randomization to TC and a statistically significant difference between maternal gut composition between the TC and nTC arms from permutation tests, we hypothesized that TC exposure affected a small proportion of taxa within the community. We identified differentially abundant taxa present in stool samples from TC vs. nTC infants and mothers by first pooling data from all three visits (Figure 4A, Supplementary table 3) and separately for each of the three visits (Figure 4B, 4C, Supplementary table 4). A greater number of taxa were significantly differentially abundant in mothers vs. infants (29 in mothers, 17 in infants; Figure 4A). Infants did not show an enrichment of specific phyla at any time point, but *Bacteroides fragilis* was persistently the most enriched species in combined and time-point analyses at 6 months (Figure 4A, 4B). Mothers showed a strong enrichment of Proteobacteria in the TC arm after the introduction of triclosan-containing toothpaste (Figure 4C, Supplementary table 4). A preliminary, unbiased quantification of triclosan resistance in the mothers using whole shotgun metagenomic sequencing of stool samples for a subset of 12 mothers in each intervention arm at 6 months was conducted. This resulted in low coverage of triclosan resistance genes; approximately 0.03% of sequenced reads mapped to triclosan antibiotic resistance genes in the Comprehensive Antibiotic Resistance Database (CARD). Unsupervised clustering by triclosan resistance gene counts was not sufficient to cluster maternal samples by intervention arm, and this may have been impacted by the limited power to detect differences between groups due to low coverage (Supplementary figure 6). Differential gene analysis from CARD showed enrichment of one antibiotic resistance gene, CfxA6, in TC households at a 10% FDR adjusted p-value threshold.

**Figure 4:**
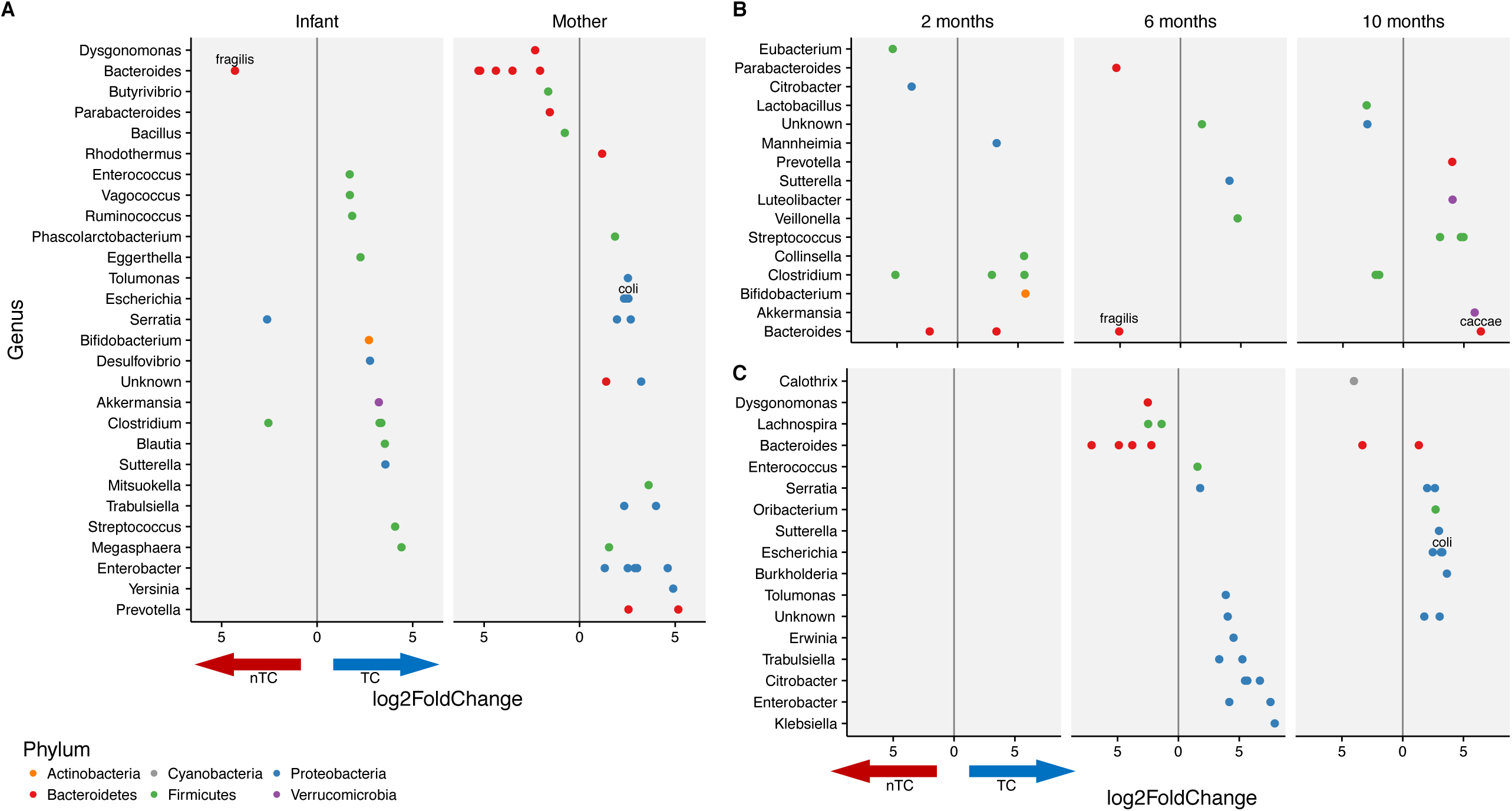
Enrichment of Proteobacteria is observed in the mothers of TC households. **(A)** Differentially abundant taxa between nTC and TC households. Values left of the grey line indicate an enrichment in nTC household and values to the right indicate an enrichment in TC households. (A) Analyses are separated by mothers and infants for all samples across the 3 time points (FDR adjusted p-value < 0.01). Differentially abundant taxa are displayed for (B) infants and (C) mothers at per visit (FDR adjusted p-value < 0.05).

## Discussion

At the time of the US FDA ruling that banned 19 antimicrobials from wash products, the extent to which TC exposure perturbed microbial populations in humans, particularly during infant development, was unknown. To test the hypothesis that exposure to TC-containing wash products induces a measurable impact on the intestinal microbiota of adults and growing infants, we assessed the stool microbiome from mothers and infants in households that had been randomized to TC or nTC wash products during the first year of the infant’s life. We observed that ongoing TC exposure from household products does not contribute to major reconstruction of either infant or adult intestinal microbiomes after approximately 10 months. TC exposure did not reduce overall gut microbial diversity in infants or mothers at any visit.

However, there are some notable trends in differential taxa with potential health implications. The most enriched species in the nTC randomized infants, *Bacteroides fragilis*, has been shown to direct maturation of the immune system(*16*) and produce anti-inflammatory polysaccharides(*17*). The most enriched organisms in the TC households at the 10-month visit were *Bacteroides caccae* in infants and *Escherichia coli* in mothers. Strain-specific triclosan resistance in *E. coli* has been described(*18*) and may explain its enrichment in TC households at the late time-point. Given that *B. caccae* was extensively enriched in the infants at the 10- month visit, it is possible that this organism also harbors interesting strategies for acquired or innate antimicrobial resistance.

In the TC arm of the study, mothers showed a strong enrichment of Proteobacteria, a phylum associated with broad spectrum antibiotic resistance. The finding that Proteobacteria were enriched in the gut microbiota is consistent with observations in fish following triclosan exposure(*5*). The emergence of Proteobacteria was associated with the introduction of triclosan-containing toothpaste after the 2-month visit. This suggests that the major intestinal exposure to triclosan is through toothpaste rather than wash products, and that personal care products not covered by the FDA ban may play a role in the expansion of antibiotic-resistant species in the intestine.

One caveat of this study is that triclosan has a relatively short half-life (24 hours) and urinary triclosan results were from only one time point; thus, it is not possible to know the cumulative dose of triclosan received by the study participants by absorption through the skin or intestine. Although mothers in the TC arm had higher levels of triclosan detected in urine than those in nTC households, the median triclosan level we detected in urine of nTC households was higher than those reported in the National Health and Nutrition Examination Surveys (NHANES) cohort from 2003-2012 (geometric mean concentration of 13.0 pg/μL, 95^th^ percentile concentration of 459.0pg/μL)(*19*). Urinary triclosan levels of mothers in TC households were approximately half of the levels following a 15 day randomization to household TC-containing products in a previously published crossover study(*9*). Some of the differences may be due to methodological variations in the protocols used for triclosan detection or differences in exposure due to geography and inconsistent product usage. Infants in TC households and nTC households often had low triclosan levels with no statistically significant difference observed between groups. Low levels are unsurprising, as infants are not using adult toothpaste in the first year of life.

Another limitation of the current study is that controlling for household TC use may not be sufficient to identify the impact of other antimicrobials on the human microbiome. We did not measure triclocarban levels directly, but exposure from external sources such as clothing and plastics (such as in children’s toys) likely occurred. Exposure to other microbicides, including any antibiotics administered throughout pregnancy and the first year, might have influenced prenatal and postnatal microbiomes in ways that cannot be experimentally controlled. However, these perturbations were present in both arms of the study. The Proteobacteria enrichment in TC mothers but not the infants suggest the small microbiota disruptions were TC dose dependent.

The impact of likely low-dose but long-term (>4 months) household product-based antimicrobial exposure on the human intestinal microbiome has not been previously described in either adults or infants during the critical phases of microbiota assembly early in life. While the impact appears to be minimal, we do identify specific taxa previously associated with anti-inflammatory properties that are enriched in nTC households, as well as other taxa previously associated with broad spectrum antibiotic resistance that are enriched in TC households. The measurable shift in the intestinal microbiome that occurs in mothers between the first and second visits of this study, which corresponds to the introduction of triclosan through toothpaste, suggests that toothpaste exposure places more selective pressure on intestinal microbial species than wash products. While a selective expansion of Proteobacteria is not known to cause various diseases, Proteobacteria expansion has been proposed as a potential diagnostic signature of dysbiosis linked to diabetes, colitis, and malnutrition(*21*). Future studies may illuminate the impact of these shifts on health-related outcomes. Evidence that antibiotic resistance develops in diverse bacterial taxa following prolonged triclosan exposure suggests that triclosan resistance may be mediated by specific genes(*22*–*24*), and that these genes may be horizontally transferred(*25*). Although we did not identify a significant enrichment of an antibiotic-resistance genes in a subset of TC-exposed mothers, future *in vitro* and potentially *in vivo* studies will be required to more thoroughly characterize the impact of TCs on antibiotic resistance in the gut microbiota. Triclosan exposure is known to play a role in allergen and food sensitization(26, 27); topical skin application of triclosan is sufficient to induce peanut sensitivity in mice(*28*). Given the high prevalence of TC exposure on the skin in this study, it will be interesting to study the impact of these wash products on the skin microbiota and related health outcomes. Despite the minimal effects of TC exposure from wash products on the gut microbial community of infants and adults, these results suggest detected taxonomic differences are associated with potential harmful effects on host physiology and highlight the need for consumer safety testing of consumer antimicrobial products on the human microbiome.

## Materials and Methods

### Study Design

Subjects in this study were recruited to participate in Stanford’s Outcomes Research in Kids (STORK), a prospective cohort study of healthy mothers and infants(*14*). Briefly, pregnant mothers were enrolled in the study at approximately 20 weeks of gestation from both Lucile Packard Children’s Hospital (Stanford, CA) and the Tully Road Clinic of Santa Clara Valley Medical Center (San Jose, CA). Enrolled mothers were additionally invited to participate in a nested, randomized intervention of TC-containing household and personal wash products to study the effects of these microbicides on illness and the development of the infant microbiome.

Participants were provided commercially available wash products (liquid and bar soap, toothpaste, dishwashing liquid) all either containing or not containing TCs. Bar soap was the only provided product that contained triclocarban in addition to triclosan. Because of concerns about potential endocrine disruption, mothers were initially not randomized to toothpaste but could continue using their preferred product during pregnancy. At the first post-delivery home visit (∼2 months post birth), either triclosan-containing or triclosan-free toothpaste was provided according to assigned arm. Supplies were replenished every four months as needed during home visits.

Household visits were conducted every 4 months to collect demographic and household information as well as stool and urine samples. Samples were stored at -80°C until processed. Automated weekly surveys on breastfeeding, diet, infant illness, including antibiotic use, were conducted, and infant medical records were referenced as available to account for antibiotic use around the time of sample collection. Of the 39 infants in this study, 34 had medical record verification of systemic antibiotic administration through the first year of life; 67.6% of infants in this study received systemic antibiotics (76.4% nTC, 60% TC) but administration did not occur within one month of sample collection in 95% of cases (104 of 109 infant samples). No antibiotic data was available for the mothers.

For intestinal microbiome analysis, we included all households with at least 5 of the expected 6 stool samples; 13 of the 39 households were missing one sample. From 13 households, one sample was missing: 5 were missing a sample from the mother (ID: 1002, 1084, 2360, 2443, 2584) and 8 from the infant (ID: 1009, 2137, 2201, 2274, 2284, 2341, 2421, 2534). To determine if the randomization to TC-containing products was sufficient to increase TC exposure, urine samples were obtained from mothers and infants at the 6-month visit to measure urine triclosan levels. This time point was chosen because it followed randomization to all products, including toothpaste.

### Urinary triclosan detection

Urinary triclosan levels (triclocarban was not assessed) were measured using liquid chromatography-mass spectrometry at the Stanford University Mass Spectrometry core facility. Urine samples (1 mL) were subjected to liquid-liquid extraction with ethyl acetate. Stable isotope labeled triclosan (13C12, 99%, Cambridge Isotope Laboratory) served as the internal standard (IS) and blank urine from subjects with no to minimal exposure to triclosan was used as sample matrix for calibration curve standards. The upper organic phase was collected and dried under a stream of nitrogen gas. Samples were reconstituted in 100uL of 20% methanol and transferred to auto-sampler vials. The LC-MS/MS analysis was performed on a TSQ Vantage triple quadrupole mass spectrometer coupled with an Accela 1250 HPLC (Thermo Fisher Scientific). Injection volume was 10 μL. Reversed phase separation was carried out on a Kinetex C18 column (50 mm × 2.1 mm ID, 2.6 um particle size, Phenomenex). Mobile Phase A was water, and mobile phase B was methanol; flow rate was 350 μL per minute. The gradient was as follows: 0 min. (20% B), 2.5 min. (98% B), 4 min. (98% B), 4.5 min (20% B) and 6 min (20% B). The mass spectrometer was operated in negative APCI mode, with selected reaction monitoring (SRM).

Three SRM transitions were used for each triclosan and triclosan IS: 250.9 > 159.1, 187.0, 214.9 and, 263.07 > 169.0, 197.9, 226.9, respectively. The calibration curve was linear from 1 to 40,000 fmol/μL, and the lower limit of quantitation (LLOQ) was around 10 fmol/μL of triclosan in extracted urine. Samples were measured in triplicate. All results were divided by 10 to account for concentration. Given the skewed distribution of triclosan levels, we used a nonparametric Mann-Whitney U test to determine triclosan level differences between intervention arms.

### DNA extraction, 16S ribosomal DNA amplification and amplicon sequencing

Samples were prepared and sequenced in two batches with approximately equal TC and nTC households per batch (29 households in batch 1, 10 households in batch 2) with identical methods. Samples were incubated for 10 minutes at 65°C before bead beating. One 20-minute round of bead-beating was performed at room temperature and samples were mixed by inversion. DNA was isolated from stool samples using the PowerSoil Isolation Kit (Mo Bio Laboratories, Inc., Carlsbad, CA) per manufacturer’s instructions. The protocol was modified to use approximately half the suggested weight (125 mg v. 250 mg) of stool per sample. Region-specific primers, which included Illumina adapter sequences and 12-base barcodes on the reverse primer, were used to amplify the V4 region of the 16S ribosomal RNA gene. Failed reactions were rerun and amplicons were cleaned using UltraClean-htp 96-well PCR Clean-Up Kit (Mo Bio Laboratories, Inc., Carlsbad, CA). Samples were quantified using Quant-iT dsDNA Assay Kit High Sensitivity (Thermo Fisher Scientific, Waltham, MA) and measured on a FLEXstation II 384 microplate reader in the Stanford High-Throughput Bioscience Center. Amplicons were then combined in equimolar ratios, ethanol precipitated and gel purified. Paired-end, 250 bp sequencing was performed on an Illumina MiSeq at the Stanford Functional Genomics Facility. A median of 47,143 (6,434-315,978) reads were sequenced per sample.

### Sequence processing and classification

We used a non-clustering method for 16S rRNA sequence classification using BaseSpace Application 16S Metagenomics v1.0 (Illumina, Inc.). Non-clustering implies amplicon reads are not grouped given a similarity score (typically 97-99%) to account for sequencing errors prior to classification. One caveat of clustering is that it reduces fine-scale variation that is biologically important. Briefly, the BaseSpace pipeline trims the 3’ ends of non-indexed reads when the quality score is less than 15. High quality reads were classified using a modified Ribosomal Database Project (RDP) Classifier (*29*) with a curated version of the Greengenes May 2013 reference taxonomy database. The original RDP classifier algorithm uses 8-base *k*-mers, however BaseSpace RDP uses 32-base *k*-mers, giving each *k*-mer more specificity for a given species. A curated version of the Greengenes May 2013 reference taxonomy filters those entries with 16S sequence length less than 1250 bp, more than 50 wobble bases, or those not classified at the genus or species level. The pipeline does not specifically check for chimeras; however, if the forward and reverse reads do not map to the same sequence in the reference database they are excluded from classification. The comparison of the BaseSpace RDP pipeline to another non-clustering sequence classification method can be found in the supplementary methods.

### Prevalence taxonomy filtering

To filter rare (i.e. noisy) taxonomically classified reads, we calculated a prevalence threshold based on taxa found in at least 7 samples. This threshold was chosen to include taxa that constitute a “core” microbiome, which suggests a taxon is persistent within at least two mothers throughout the study or a taxon during development is found in at least 20% of infants at one visit. This filtering resulted in the inclusion of 1115 taxa from a pool of 1892 taxa.

### Metagenomic characterization of antibiotic resistance profiles

DNA archived from the previously described stool extraction was processed for shotgun DNA sequencing using the Nextera XT DNA Library Preparation Kit (Illumina Inc.) per manufacturer’s instructions on a subset of 24 maternal samples from the 6-month visit. Paired-end, 101 bp sequencing was performed on an Illumina HiSeq 4000 at the Stanford Sequencing Service Center with an average of 21,660,032 (12,825,854–31,271,994) reads per sample.

Low quality read ends with a Phred score less than 20 were trimmed using TrimGalore v. 0.4.1 (http://www.bioinformatics.babraham.ac.uk/proiects/trim_galore/), and PCR duplicates were removed using Super-Deduper v. 1.40 (http://dstreett.github.io/Super-Deduper/). High quality reads were then aligned to the Comprehensive Antibiotic Resistance Database (CARD)(*30*) using Burrows-Wheeler Aligner v. 0.7.10 (http://bio-bwa.sourceforge.net/). A median of 25,064 (8200-72,464) reads mapped to CARD per sample, with a median mapping percent of 0.11% for both intervention arms after adjusting for number of sequenced reads. Genes known to harbor triclosan resistance (reviewed in (*31*) mapped a median of 5,476 (763-22,205) reads per sample. TC samples had a median of 7,043 mapped reads (relative % adjusted for sequencing depth: 0.030%) compared for 5,330 reads (relative % adjusted for sequencing depth: 0.027%) for nTC samples. Euclidean distance was calculated between samples, then clustered with a hierarchical agglomeration method using base R functions.

### Statistical Analyses

Analyses were performed in ‘R’ v. 3.2.4 (http://www.R-project.org) with accompanying packages on non-rarified unique taxonomic classifications(*32*). Principle coordinate analyses (PCoA) were performed using non-metric Bray-Curtis dissimilarity for combined analyses using ‘phyloseq’ v. 1.14(*33*). Levels of triclosan in household products are intended to inhibit bacterial growth rather than kill bacteria(*34*); Because we hypothesized that the suppression of growth would alter taxa abundances rather than the presence/absence of taxa, Bray-Curtis, a non-phylogeny based method that takes abundance into account, was chosen for PCoA. Sample distances for maternal-only analyses were calculated using the Canberra distance, which is best used for centroid type patterns since maternal points on the overall PCA were centralized (Figure 1).

Alpha diversity measured by the Shannon diversity index was calculated using ‘vegan’ v. 2.4(*35*). Nonparametric Mann-Whitney U tests were used to determine statistical differences in diversity between intervention arms at each time point given the low sample sizes per comparison. To statistically test treatment effects on the homogeneity of microbial community composition, we performed permutational multivariate analysis of variance (PERMANOVA) analyses on distance metrics with ‘vegan’ v. 2.4; 1000 permutations were performed for mother and infant groups, stratified by the visits to account for infant development and maternal toothpaste introduction after the first visit as confounders. Differential taxa abundance and gene analyses were performed using ‘DESeq2’ v. 1.10(*36*). A 1% FDR adjusted p-value threshold was selected for the treatment comparisons across mothers and infants pooling all three visits. This threshold is relaxed to 5% for comparisons at each time visit given lowered power from smaller sample size.

## Acknowledgements

The authors thank Karolina Krasinska and the Vincent Coates Foundation Mass Spectrometry Laboratory, and the Stanford University Mass Spectrometry Laboratory (http://mass-spec.stanford.edu) for assistance developing and running the triclosan assays, and Susan Holmes and Eli Moss for helpful discussions regarding the analyses.

## Author Contributions

J.P. and C.L. conducted the cohort study and nested intervention from which the specimens were collected and oversaw collection and coordination of all samples and metadata. T.D.H. coordinated processing of stool and urine samples and extracted and amplified DNA from the stool samples. E.T. prepared the shotgun sequencing libraries. A.S.B., J.V.R. designed the data analysis; J.V.R. performed the computational and statistical analysis; J.V.R., A.S.B., J.P. participated in data analysis and manuscript preparation; all authors edited the manuscript.

---

**Supplementary table 1: Summary of number of sequenced and classified reads per sample.**

Classified reads reported have been filtered to remove rare, i.e. noisy, taxa.

**Supplementary table 2: Urinary triclosan levels at 6 months.**

Triclosan was measured using liquid chromatography-mass spectrometry in urine samples collected at the second visit (∼6 months). Summaries are provided as the median (range) of TC molecules in a picogram/microliter (pg/μL).

**Supplementary table 3: Differentially abundant taxa for infants and mothers aggregated across 3 visits.**

Differentially abundant taxa using DESeq2 (adjusted p-value < 0.01) with 1115 unique taxonomic classifications.

**Supplementary table 4: Differentially abundant taxa in TC and nTC households by group and visit.**

Differentially abundant taxa were determined using DESeq2 (adjusted p < 0.05) with 1115 unique taxonomic classifications. No species were differentially abundant for mothers at 2 months.

**Supplementary figure 1: Distribution of sample collection relative to infant birth.**

Households for each visit time point are sampled approximately within one month, and are balanced between treatments in the study.

**Supplementary figure 2: DADA2 and BaseSpace RDP algorithms comparably capture microbiome variance.**

Principal coordinates analyses of Bray-Curtis dissimilarity of “core” or “developmental” taxa show that gut communities cluster by mothers and infants. **(A)** DADA2 and **(B)** BaseSpace RDP are comparable in first and second axis variance.

**Supplementary figure 3: Individuals in mother and infant groups have similar relative microbiome compositions at the phylum level throughout the first year of life independent of TC exposure.**

**(A)** TC households (n=17) and **(B)** nTC (n=22) households have stable relative abundance of species at 2, 6, and 10 month visits. Analyses are complete, with only 13 missing samples total out of 234 proposed for the study (94% inclusion). Phyla present in less than 2% abundance are condensed for visual clarity.

**Supplementary figure 4: Infant intestinal microbiome variability is not defined by known external factors by 2 months of age (n=34).**

PCoA with Bray-Curtis dissimilarity of the infants at 2 months of age suggests factors known to impact the microbiome, such as **(A)** birth method and **(B)** formula, **(C)** maternal ethnicity, and **(D)** pets in the household are not the main drivers of microbiome variance in infants at 2 months of age.

**Supplementary figure 5: Maternal samples within households are more similar than between households.**

**(A)** Hierarchical clustering of Canberra distances shows that maternal samples from a given individual throughout the first year of life are more self-similar than other household mothers at the same visit. The colors are representative of a single household, labeled by household and visit (B1 = 2 months, B2 = 6 months, B3 = 10 months), and node colors reflect TC grouping (red=nTC, blue = TC). **(B)** PCoA of Canberra distances shows 9.3% of the variance is explained by the first two axes and the absence of TC treatment clustering for any visit among mothers. **(C)** PCoA projection variability between visits occurs similarly between TC and nTC households when compared to the initial 2-month visit.

**Supplementary figure 6: Triclosan resistance gene abundances do not distinguish TC and nTC maternal samples following 6 months of exposure.**

Metagenomic reads were aligned to the CARD Database. Known triclosan resistance genes had a median mapping percent of ∼0.03% for both intervention arms after adjusting for number of sequenced reads. Euclidean distance was calculated between samples, then clustered with a hierarchical agglomeration method. Mothers of both treatment arms have a low number of reads in triclosan resistance genes (∼6,000 reads) suggesting we limited power to detect difference with this approach and require targeted sequencing or culturing studies to further understand resistance profiles in the stool.

